# Hallmarks of glucocorticoid receptor condensates involvement in transcription regulation

**DOI:** 10.1101/2024.11.14.623561

**Authors:** Belén Benítez, Martin Stortz, María Cecilia De Rossi, Diego M. Presman, Valeria Levi

## Abstract

Several proteins necessary for mRNA production concentrate in intranuclear condensates, which are proposed to affect transcriptional output. The glucocorticoid receptor (GR) is a ligand-activated transcription factor that regulates the expression of hundreds of genes relevant to many physiological and pathological processes. As with all members of the steroid receptor family, GR forms condensates of unknown function. Here, we examine whether GR condensates are involved in transcription regulation using Airyscan super-resolution microscopy and nano-antibodies targeting initiation and elongating states of RNA polymerase II (Pol2). We observed subpopulations of GR condensates colocalizing with initiating and, surprisingly, elongating Pol2 foci. The analysis of GR mutants with different transcriptional outputs suggests a correlation between condensate formation capability and transcription initiation. Moreover, the number of GR molecules within initiation and elongation condensates appears to be linked to transcriptional activity. Taken together, our data suggests an involvement of GR condensates in transcription initiation and elongation.

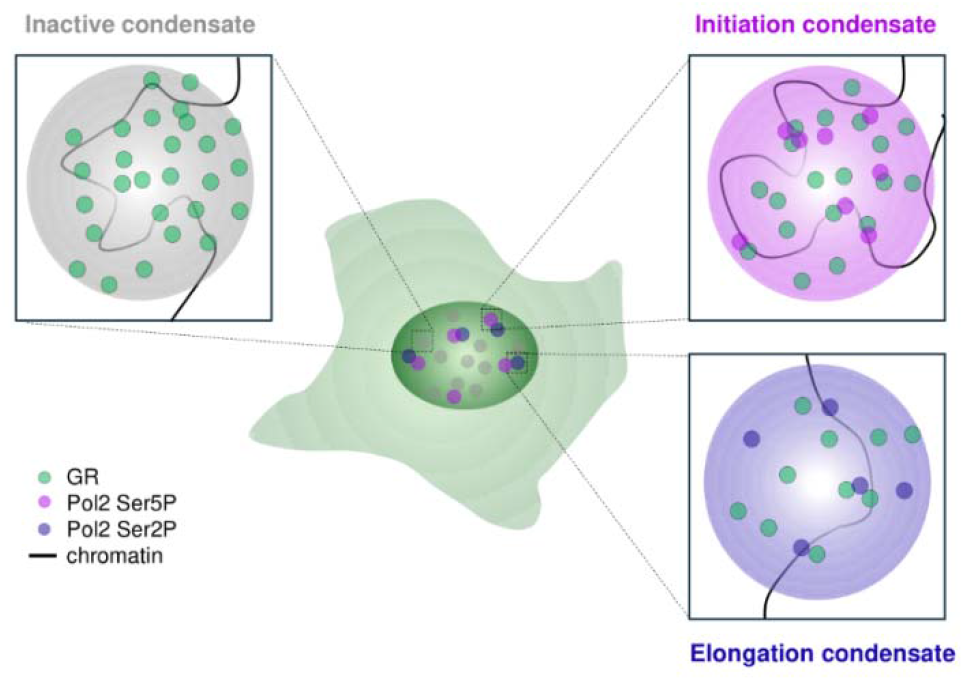

## Introduction

The nucleus of eukaryotic cells is a crowded place, wherein chromatin is organized at different spatial length scales, from the 10-nm chromatin fiber up to micrometer-sized chromosome territories [1, 2]. How do transcriptional regulators access the genome efficiently to control the expression of their target genes is a matter of continuous debate.

The observation of diverse membrane-less compartments concentrating molecules related to specific nuclear activities led to the idea of a hierarchical organization in which nuclear processes are spatially confined within specific regions in the nucleus [3]. This model gained visibility with the proposal of liquid-liquid phase-separation (LLPS) as the underlying process promoting the formation of many nuclear membrane-less structures now referred to as liquid condensates [4]. Nevertheless, it has been proposed that weak and transient interactions between multivalent biomolecules could also account for many of the properties attributed to liquid condensates, without the need to invoke LLPS [5].

Independently of the underlying process, it is now widely accepted that many transcription factors (TFs) concentrate in distinct nuclear foci which are referred to as ‘transcriptional condensates’ because they also accumulate other transcriptional players such as RNA polymerase II (Pol2), the Mediator complex, and coregulators [6]. Initial studies claimed that these condensates promote transcription by concentrating transcription-activating molecules near gene promoters [7]. This initial view evolved as the link between condensates and transcriptional regulation has proven to be more complex, with transcriptional condensates having either activating, inhibitory, or neutral roles on gene transcription in different conditions [reviewed in [6]].

In recent years, it has been proposed that the different steps of the transcription process, including the formation of pre-initiation complexes, elongation, and RNA splicing, are confined into separated condensates [8, 9]. This physical segregation may be due to the selective partitioning of biomolecules driven by, for example, the charge patterns of IDRs [10]. In line with this model, the phosphorylation of the C-terminal domain (CTD) of Pol2 promotes its dissociation from condensates enriched in molecules related to transcription initiation and its incorporation into those involved in RNA processing [11], also suggesting that condensates might play a role in the progression of transcription. In a similar direction, the Young group proposed that transcriptional condensates are subject to a feedback mechanism of transcriptional regulation, as low levels of nascent RNA favor the formation of transcriptional condensates through electrostatic interactions whereas relatively higher levels of RNA produced during the bursting events contribute to their dissolution [12].

Steroid receptors (SRs) are a family of TFs that regulate the expression of thousands of genes upon binding to steroid hormones and play important roles in both physiological and pathological processes [13]. This family includes the glucocorticoid, androgen, progesterone, estrogen, and mineralocorticoid receptors, which share the property of forming nuclear condensates when activated by hormones [14]. Even though these condensates were first described decades ago [15], their biological function remains elusive. The glucocorticoid receptor (GR), a member of the SR family, regulates a plethora of physiological functions ranging from metabolism to the immune system response [16]. The inactive receptor distributes in the cytoplasm, and upon ligand activation, it translocates into the nucleus forming multiple condensates [17, 18]. We and others have previously shown that GR foci present many properties compatible with liquid condensates, require a chromatin scaffold and incorporate other transcriptional players [19-21]. These observations suggest their involvement in transcription regulation, however direct evidence to support this hypothesis is still lacking.

Here, we analyze if GR condensates participate in transcription using Airyscan microscopy to improve the spatial resolution [22], and nano-antibodies designed to target the initiating and elongating Pol2 phosphorylated forms [23, 24], allowing their visualization in live cells. By using these tools, we observed populations of GR condensates engaged in transcription initiation and elongation. We also studied a possible link between GR condensates and transcription regulation using GR mutants with different transcriptional behavior. Our results suggest a correlation between the ability of the receptor variants to form condensates and its involvement in transcription initiation while the number of receptor copies included in initiation and elongation condensates appears to be linked to the receptor’s transcriptional activity.

## Results

### Subpopulations of GR condensates colocalize with active forms of RNA polymerase II

Upon ligand activation, the glucocorticoid receptor (GR) forms nuclear condensates (Fig. 1A), with some of their properties recently characterized by us and others to be compatible with liquid condensates [20, 21]. However, the role (if any) of GR condensates in transcriptional regulation remains elusive [6]. Most condensates formed by transcriptional-related molecules, including those of GR, are sub-diffraction-sized structures, and therefore, their analysis is limited by the optical resolution of conventional fluorescence microscopy setups. To visualize these condensates with improved spatial resolution compared to previous studies, we used Airyscan microscopy [22] for all imaging experiments. Figure 1A shows the comparison between standard confocal and Airyscan super-resolution modes when imaging condensates of EGFP-tagged GR transiently expressed in murine mammary adenocarcinoma cells wherein endogenous GR has been knocked out [D4 cells, [25]]. The images obtained after stimulation with the synthetic GR agonist dexamethasone (Dex) showed a noticeable improvement in spatial resolution and condensates detection.

**Figure 1.**
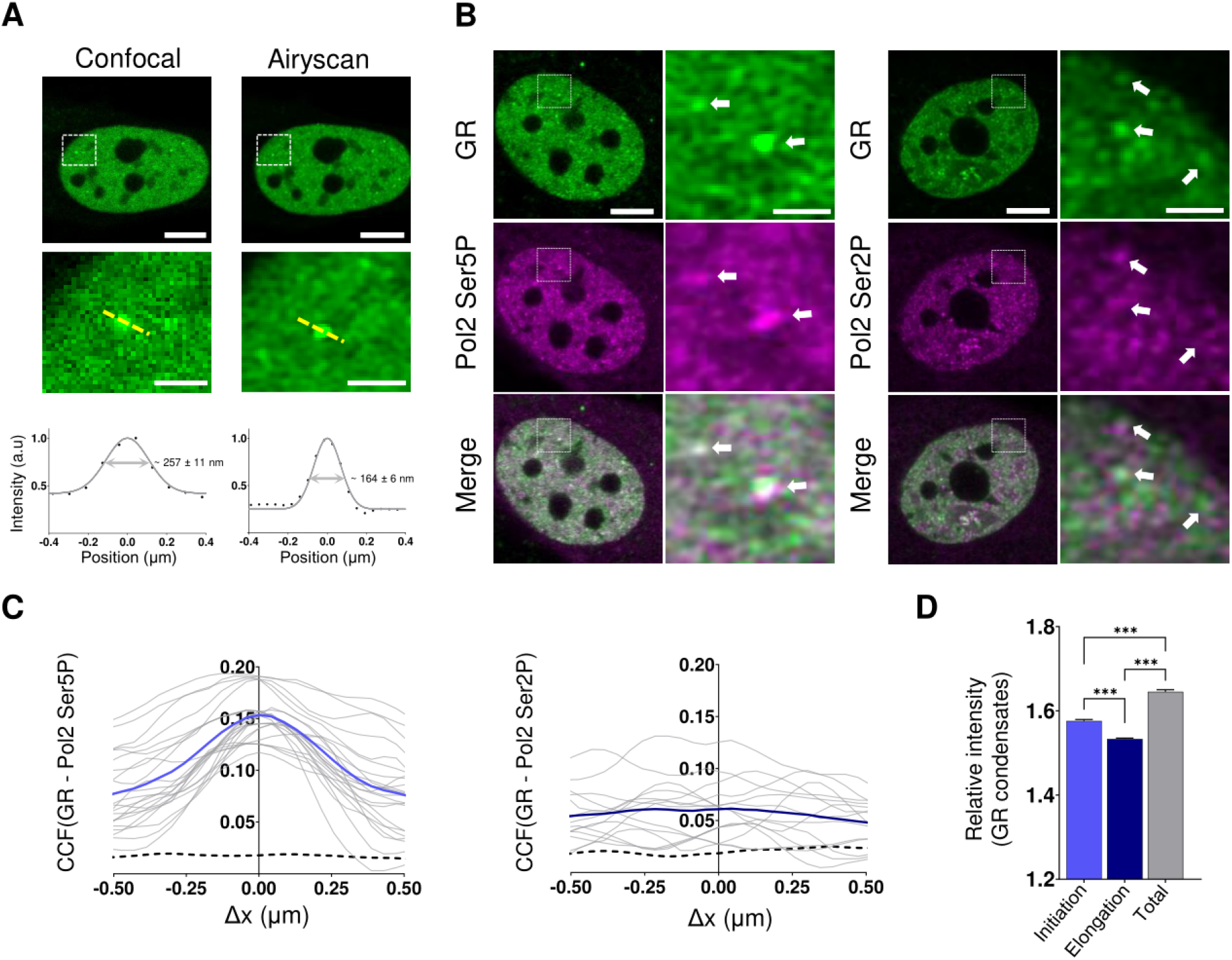
GR condensates spatially correlates with active Pol2 foci. A. Representative confocal and Airyscan images of D4 cells transiently expressing EGFP-GR (green), (left panels). Scale bar, 5 µm. Zoom-in images of a GR condensate (white square) and its intensity profile indicated by the yellow dotted line (right panels). Scale bar, 1 µm. The full width at half maximum (arrow) was determined by a Gaussian fitting. B. Representative Airyscan images of D4-Halo-GR cells labeled with JF549 (green) and transiently expressing nano-antibodies fused to GFP-tagged Pol2 Ser5P or Pol2 Ser2P (magenta). Scale bar, 5 µm. Zoom-in images of the region delimited by the white squares show the overlap of GR condensates and initiation and elongation Pol2 foci (arrows). Scale bar, 1 µm. C. Colocalization analysis of GR condensates and GFP-tagged nano-antibodies against Pol2 Ser5P or Pol2 Ser2P foci (top and bottom panels, respectively). The average CCF curve was calculated for each experimental condition (blue line) and compared to that expected for uncorrelated events (dashed line). D. Relative intensities of the total GR condensates and of those colocalizing with Pol2 Ser5P (initiation) or Pol2 Ser2P (elongation) foci. Data is expressed as means ± SEM. Data information: Data sets are representative of at least three independent experiments. The number of cells (n) was: (A) n_GFP-GR confocal_ =15 and n_GFP-GR Airyscan_ =15; (C) n_Halo-GR/GFP-Pol2 Ser5P_ = 23; (D) n_Halo-GR/GFP-Pol2 Ser2P_ = 14. Statistical analysis was performed by Man Whitney’s test. ns = not significant; * p < 0.05; ** p < 0.01 and *** p < 0.001.

If GR condensates are involved in transcriptional activation, they should be found colocalizing or in close physical proximity to the active transcriptional machinery. RNA polymerase II (Pol2) is subject to several post-translational modifications that are intimately related to its activity. The phosphorylation of the catalytic subunit within its long heptapeptide repeat located in the CTD defines its transcriptional status: Ser5 phosphorylation (Ser5P) marks initiating Pol2 while Ser2 phosphorylation (Ser2P) is a hallmark of an elongating polymerase [26]. The Kimura lab has recently developed genetically encoded nano-antibodies fused to GFP against each of these active Pol2 forms [23, 24], allowing the observation of both initiating and elongating endogenous Pol2 in live cells.

To test whether there is a spatial correlation between GR condensates and the active forms of Pol2, we transiently expressed each GFP-tagged antibody against Pol2 Ser2P or Pol2 Ser5P in D4 cells stably expressing HaloTag-tagged GR (D4-HaloGR). As previously reported [11], both initiating and elongating Pol2 molecules distribute in foci within the nucleus (Fig. 1B).

Representative live-cell images acquired after GR activation with Dex for at least 30 min, show that, in some cases, Pol2 Ser5P and Pol2 Ser2P foci partially overlapped with GR condensates (Fig. 1B). In line with this qualitative observation, an image correlation analysis (Fig. S1A) revealed a zero-centered positive spatial cross-correlation function (CCF) between GR condensates and foci of Pol2 Ser5P (Fig. 1C). We also detected a small but positive cross-correlation with Pol2 Ser2P foci suggesting some colocalization between GR condensates and elongating foci (Fig. 1C). We measured the fraction of the total area of GR condensates overlapping with phosphorylated Pol2 foci (f_GR_,, Fig. S1B) and confirmed that the degree of overlap with both forms of Pol2 was significantly higher than the expected for unrelated foci distributions (Table 1). These data also suggest that ∼19% of the area of GR condensates colocalizes with initiating Pol2 (Ser5P) whereas ∼10% overlaps with elongating Pol2 (Ser2P), suggesting that only a fraction of GR condensates would be actively engaged in transcription while most of them remain inactive within the imaging time window (18.4 s).

**Table 1.** Proportion of total area of GR condensates that colocalize with Pol2 foci.

| Receptor | $f_{GR}$ (%) ($^{\Phi}$) | | $N_{Condensates} / N_{Cell}$ | |
| --- | --- | --- | --- | --- |
|  | Pol2 Ser5P | Pol2 Ser2P | Pol2 Ser5P | Pol2 Ser2P |
| GRwt | 19.1 ± 3.3 | 10.4 ± 1.5 | 690 / 23 | 352 / 14 |
| GRdim | 19.3 ± 1.2 | 6.1 ± 0.6 | 205 / 31 | 84 / 27 |
| GRtetra | 15 ± 1 | 6.3 ± 0.9 | 1283 / 25 | 896 / 25 |

( $^{\Phi}$ ) Data is expressed as means ± SEM

Collectively, our data suggests the existence of at least three subpopulations of GR condensates: those that colocalize with Pol2 Ser5P (herein referred to as ‘initiation condensates’), those colocalizing with Pol2 Ser2P (‘elongation condensates’), and the ones that participate in neither of these processes during the snapshot time window (inactive condensates). Interestingly, the mean intensity of elongating condensates was lower than that of initiation condensates (Fig. 1D). On the other hand, both GR condensate subpopulations present lower mean intensities compared to all nuclear GR condensates (Fig. 1D), suggesting that inactive condensates include more GR molecules per condensate than the active ones.

The D4 adenocarcinoma cells have a tandem array of ∼200 copies of the GR-responsive promoter MMTV driving *Ras* gene expression (MMTV-array) that can be visualized in ligand-stimulated cells as a bright region in the nucleus due to the recruitment of fluorescently labeled GR to its specific targets [27] (Fig. S2A). Even though the MMTV-array presents a complex, non-simultaneous transcriptional behavior wherein not all [28], the results obtained at GR condensates mirrored those at the MMTV-array, as we observed partial colocalization with Pol2 Ser5P and, to a lesser extent, with Pol2 Ser2P foci (Fig. S2B). Hence, these markers are suitable for detecting transcription initiation and elongation, respectively.

The colocalization of GR condensates with both, initiating and elongating Pol2 could just be the result of the dynamic, progressive transcriptional status of the active genes within each GR condensate. However, it has been proposed that initiating and elongating Pol2 molecules differ in their spatial organization [11, 23]. Even though we cannot simultaneously label Pol2 Ser5P and Pol2 Ser2P to further explore the subnuclear compartmentalization of these two forms of Pol2, we independently analyzed the colocalization of each of them with mCherry-tagged BRD4 (Fig. 2A and B), a transcriptional coactivator that forms part of the initiation complex [29]. In line with the hypothesis of a physical separation between initiating and elongating condensates, we observed that BRD4 foci present a non-shifted, spatial correlation with Pol2 Ser5P foci (Fig. 2C). In contrast, the CCF analysis with Pol2 Ser2P foci presents a ∼0.3 µm shift (Fig. 2D). These results strongly suggest that at least some transcription initiation and elongation events occur in neighboring compartments physically separated in our cell system.

**Figure 2.**
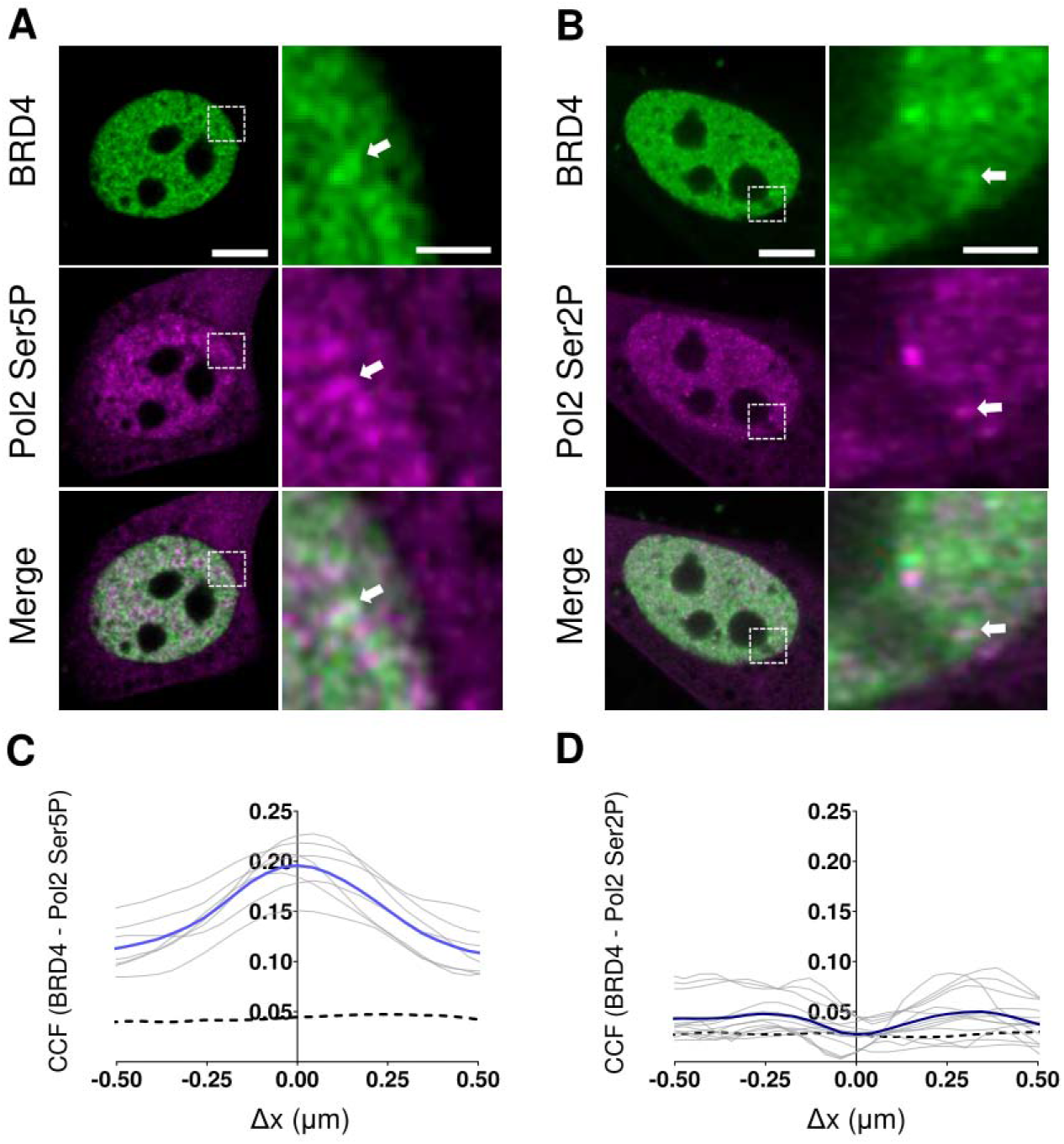
Differential spatial distribution of BRD4 and Pol2 foci. A,B. Representative Airyscan images of D4 cells co-expressing mCherry-BRD4 (green) and GFP-tagged nano-antibodies against GFP-Pol2 Ser5P (A, magenta) or GFP-Pol2 Ser2P (B, magenta). Scale bar, 5 µm. Zoom-in images of the region delimited by the white squares show the relative position of BRD4 with respect to Pol2 foci (arrows). Scale bar, 1 µm. C,D. Colocalization of BRD4 and Pol2 Ser5P (C) or Pol2 Ser2P (D) foci. The average CCF curve for each experimental condition (blue line) and that expected for uncorrelated events (black line) are shown. Data information: Data sets are representative of at least two independent experiments. The number of cells (n) was: (C) n_mCherry-BRD4/GFP-Pol2 Ser5P_ = 10 and (D) n_mCherry-BRD4/GFP-Pol2 Ser2P_= 13, respectively.

Taken together, our data indicates that subpopulations of GR condensates engage in the initiation and elongation steps of the transcription process with higher participation in the former, supporting a role for GR condensates in transcriptional regulation.

### The concentration of GR in condensates can be regulated by transcriptional modulators

As shown above (Fig. 1D), the relative concentration of GR molecules per condensate differs between initiating and elongating condensates. Therefore, further insights can potentially be obtained by analyzing the intensity distribution of all GR condensates. This distribution is positively skewed (Fig. 3A and Table S1), indicating the existence of a subpopulation of brighter condensates that probably require IDRs-dependent interactions as the intensity distribution of a truncated GR version lacking the disordered N-terminal domain (GR407C mutant) lacks this brighter subpopulation (Fig. 3A). As the dimmer GR condensates are less likely to be detected, especially those with intensities close to the intensity threshold (see Methods), their contribution to the left branch of the distribution could be underestimated. Therefore, we fitted the right branch with an empirical, exponential-decay function to further quantify the heterogeneous population of condensates (Fig. 3A and Table S1).

**Figure 3.**
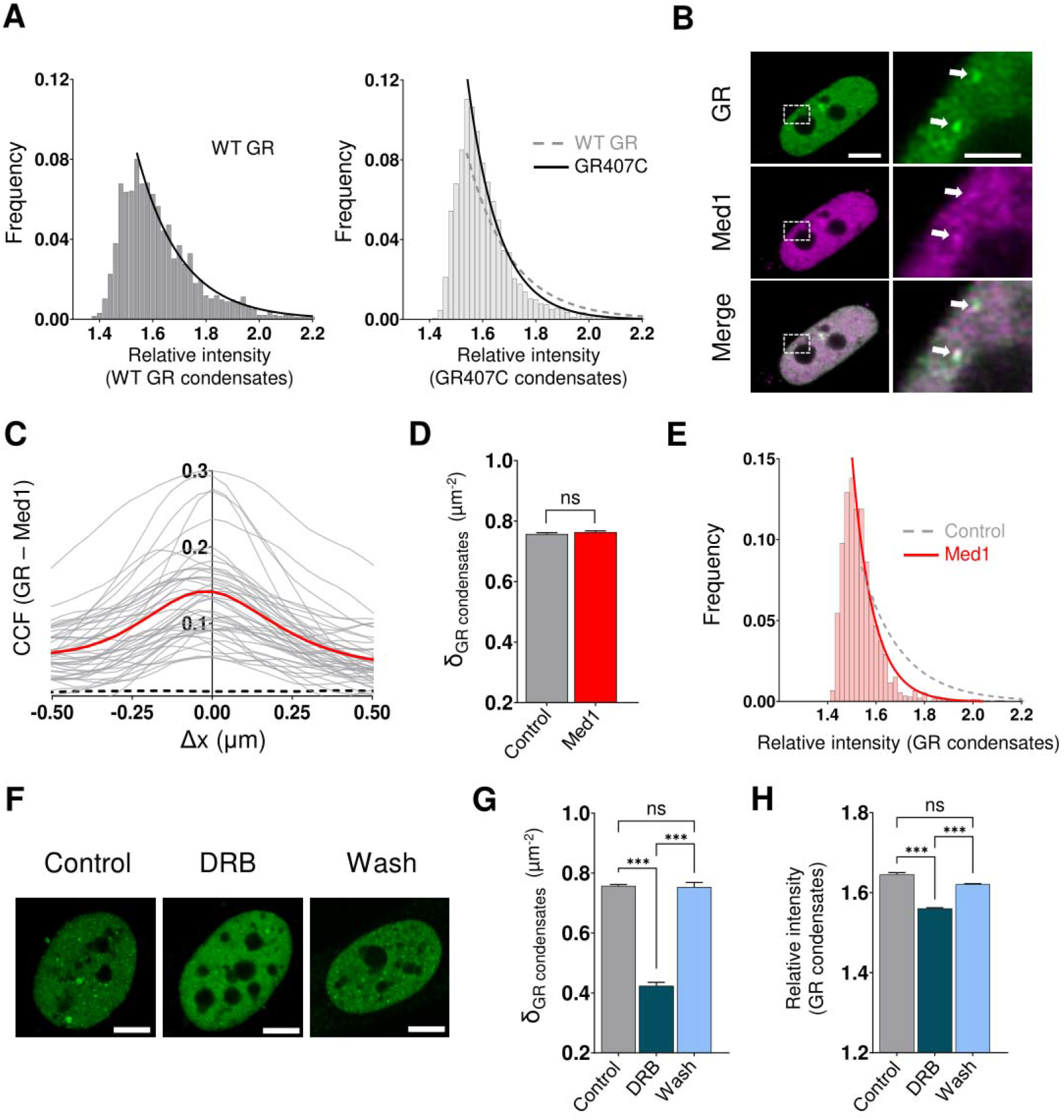
GR condensates are sensitive to transcriptional modulators. A. Intensity distributions of WT GR and GR407C condensates registered in D4 cells. The right tail of the distributions was fitted with an exponential-decay function (black line) obtaining the parameters reported in Table S1.B. Representative Airyscan images of D4 cells transiently expressing EGFP-GR (green) and Halo-Med1 labeled with JF549 (magenta). Scale bar, 5 µm. Zoom-in images of the region delimited by the white squares show the overlap of GR condensates and Med1 foci (arrows). Scale bar, 1 µm. C. Colocalization of GR condensates and Med1 foci. The average CCF curve (red line) and that expected for uncorrelated events (black line) are shown. D. Density of GR condensates in D4 cells expressing EGFP-GR (control) or EGFP-GR and Halo-Med1 (Med1) labeled with JF549. Data is expressed as means ± SEM. E. Intensity distribution of GR condensates in D4 cells co-expressing EGFP-GR and Halo-Med1 labeled with JF549. The right branch of the distribution was fitted with an exponential-decay function (red line) obtaining the parameters reported in Table S1. To facilitate comparison, the fitting curve obtained for GR in the absence of Med1 is also shown (grey dotted line). F. Representative Airyscan images of Dex-stimulated D4 cells expressing EGFP-GR (green) registered without DRB (control condition), after incubation with DRB (DRB) and after drug removal (Wash). Scale bar, 5 µm. G. Density of GR condensates in D4 cells in control condition, after DRB incubation (DRB) and after drug removal (Wash). Data is expressed as means ± SEM. H. Relative intensity of GR condensates in D4 cells in control condition, after DRB incubation (DRB) and after drug removal (Wash). Data is expressed as means ± SEM. Data information: Data sets are representative of at least three independent experiments. The number of cells (n) was: (A) n_EGFP-GR_ = 46 and n_EGFP-GR407C_ = 37; (C) n_EGFP-GR/Halo-Med1_ = 46; (D-E) n_EGFP-GR_ = 46 and n_EGFP-GR/Halo-Med1_ = 46; (G-H) n_EGFP-GR_ = 46 and n_EGFP-GR+DRB_ = 70. Statistical analysis was performed by Student’s t-test or unpaired t-test with Welch’s correction. ns = not significant; * p < 0.05; ** p < 0.01 and *** p < 0.001.

To get insights into the molecular processes underlying the heterogeneous intensity distribution of GR condensates, we next analyzed if the interactions with molecules relevant to transcription can influence this distribution. Mediator is a multi-subunit complex involved in transcription regulation [30] that forms liquid condensates in association with chromatin and other transcriptional players [31]. Med1, a subunit of the Mediator complex, acts as a GR coactivator in a ligand-dependent manner [32] and it incorporates into GR condensates in U2OS cells [20]. As expected, co-expression of EGFP-GR and Halo-Med1 in D4 cells revealed partial colocalization between GR and Med1 condensates (Fig. 3B and C). Although the overexpression of Med1 did not significantly affect the density of GR condensates (Fig. 3D), its intensity distribution shifted towards lower intensity values (Fig. 3E and Table S1). The loss in the distribution’s tail suggests that the incorporation of Med1 into GR condensates promotes the release of GR molecules to the nucleoplasm. Similarly, the expression of Halo-Med1 caused the dissociation of GR molecules from the MMTV-array (Fig. S2C). Thus, the Mediator subunit does not passively incorporate into an already-formed GR condensate, but it triggers a reconfiguration in the network of molecular interactions preexistent at the condensate, leading to a new composition with a lower amount of GR molecules in condensates.

To further explore the relationship between transcriptional activity and the intensity distribution of GR condensates, we used the reversible transcriptional inhibitor 5,6-dichloro-1-beta-D-ribofuranosylbenzimidazole (DRB). This compound blocks the transition of Pol2 to productive elongation [33], but it does not prevent the already started elongation events from finishing transcription [34]. We verified that 100 µM of DRB for 30 min reduced both the density and mean intensity of Pol2 Ser2P foci while it did not affect Pol2 initiation foci (Fig. S3A-C). Surprisingly, similar treatment in Dex-stimulated GFPGR-D4 cells reduced GR intensity at the MMTV array (Fig. S2C), indicating that transcriptional elongation appears somehow involved in GR recruitment at specific response elements within the array. In the same direction, DRB treatment produced the dissolution of multiple GR condensates (Fig. 3F-H). These observations are consistent with the idea that at least a subpopulation of GR condensates engages in active transcription. Furthermore, it also suggests that transcriptional activity itself contributes to the maintenance of some GR condensates.

DRB treatment also affected the intensity distribution of GR condensates by skewing it towards lower intensity values (Fig. S3D and Table S1), in line with the observations at the MMTV array [35] (Fig. S2C) and, once again, highlighting how active transcription also modulates GR condensates’ composition. The comparative analysis of the intensity distributions, the mean intensity and foci density between non-treated and DRB-treated cells shows a reversible loss of high-intensity foci (those with relative intensity > 1.7) after inhibiting elongation (Fig. 3F-H) which represent only ∼ 27 % of the total GR condensate population. Thus, this does not rule out that those dimmer GR condensates engaged in elongation might also be compromised by DRB as this drug affects ∼45% of the total GR condensates (Fig. 3G), which is higher than the entire population of brighter condensates. The variability in elongation foci’s intensity could be partly related to the proposed scaffolding properties of nascent RNA in transcriptional condensates, producing changes in the electrostatic balance and the composition of condensates as elongation proceeds [12].

### GR mutants with different transcriptional activities present distinct abilities to engage in initiation and elongation condensates

To further explore the functional link between GR condensates and gene expression regulation, we analyzed the behavior of two GR mutants that form transcriptional condensates but present different transcriptional profiles. Specifically, we selected the GR-P481R mutant, referred to as GRtetra, which is constitutively tetrameric in contrast to the wild-type (WT) GR that forms dimers upon hormone stimulation and only tetramerizes when bound to DNA targets [27]. Compared to the WT GR, GRtetra binds more specific chromatin sites and regulates more genes [36]. On the other side of the transcriptional spectrum, we chose the GR-A465T mutant (known as GRdim) that binds ∼2/3 of the GR binding sites genome-wide [25, 37] but only presents a modest capability of activating the transcription of target genes [25].

In line with previous reports [27, 38], we observed that GRtetra and GRdim form a large number of condensates upon stimulation with Dex both to a greater extent than the WT receptor (Fig. S4A and B), indicating that the ability of a GR mutant to form condensates is not sufficient to predict its transcriptional output. However, the intensity distribution of the condensates formed by these mutants showed almost opposed behaviors when compared to GR (Fig. S4C and Table S1). Specifically, GRtetra condensates’ intensities spanned a slightly larger intensity range, suggesting that the exacerbated multivalency of the tetrameric receptor improves the ability of the receptor to interact with molecular partners stabilizing larger condensates. In contrast, GRdim presented a lower contribution of brighter condensates to its intensity distribution in comparison to GR (Fig. S4C and Table S1), suggesting some impairment in this mutant to interact with certain components of the condensates.

To assess how these GR mutants associate with Pol2 foci, we co-transfected D4 cells with HaloTag-GRdim or HaloTag-GRtetra and either GFP-tagged nano-antibody against Pol2 Ser2P or Pol2 Ser5P (Fig. 4A). We first analyzed the spatial relationship between the condensates formed by GRdim or GRtetra with the transcription initiation/elongation foci. CCF analysis revealed colocalization between GRtetra and GRdim condensates with both Pol2 Ser5P and Pol2 Ser2P foci (Fig. 4B) although the shape of the curves suggests a different spatial relationship. While the CCF functions of GRtetra show peaks at zero-shift, indicating that the tetrameric receptor engages with both initiation and elongation foci, only the correlation curve of GRdim with Pol2 Ser5P foci shows a similar peak. In contrast, the CCF of GRdim condensates with Pol2 Ser2P foci is flatter and presents small local maxima at ∼ 0.3 μm shift, reminiscent of the behavior observed for BRD4 with elongation foci (Fig. 2D) as well. These results suggest a comparatively lower involvement of GRdim in Pol2 elongation foci, compared to GRtetra and WT GR. In addition, the analyses of the relative area of GRtetra and GRdim condensates overlapping with the phosphorylated Pol2 foci (f_GR_) revealed that most of the mutants’ condensates remain inactive during the studied time window as observed for the WT receptor (Table 1).

**Figure 4.**
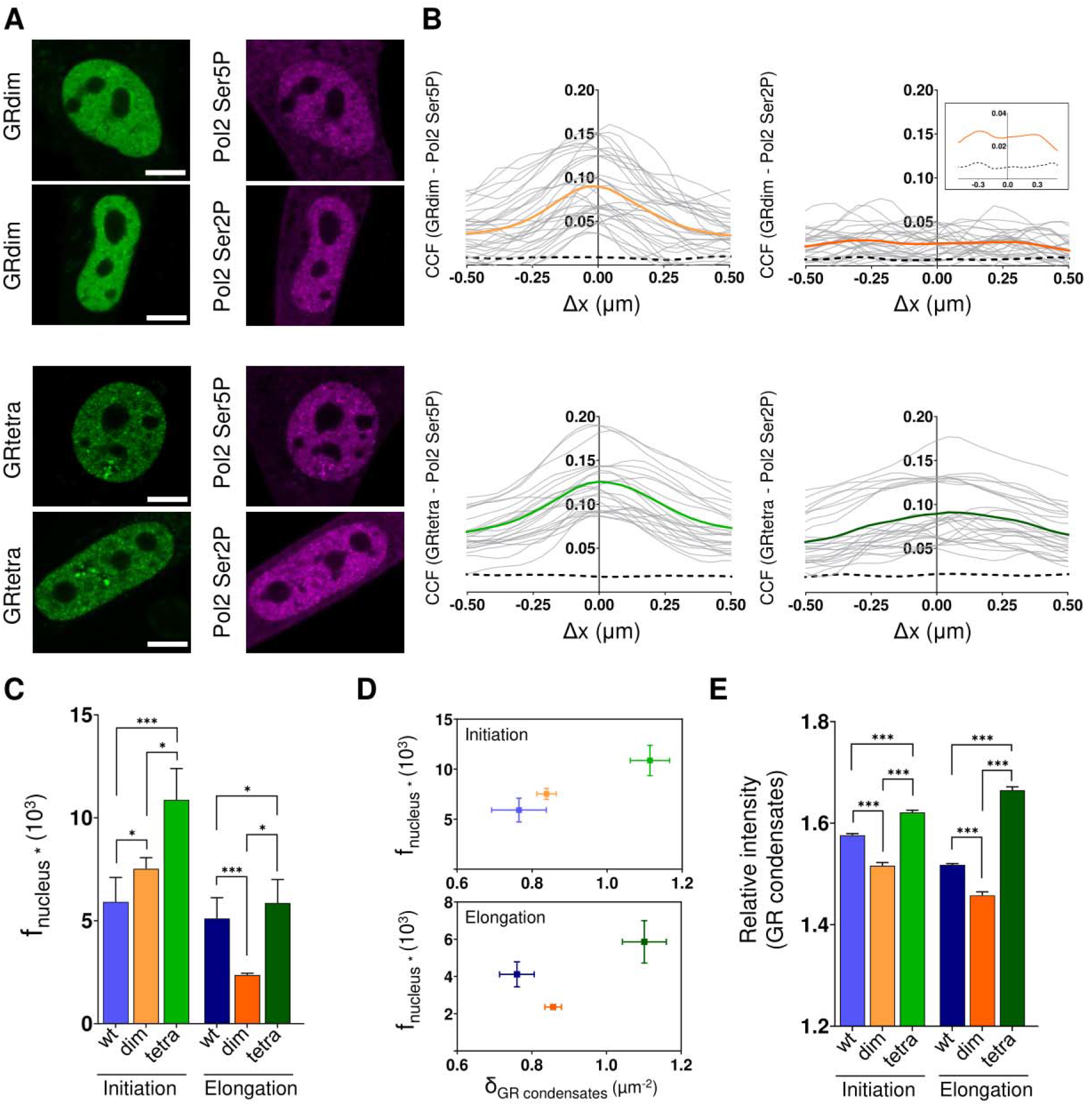
GR mutants associate with initiation and elongation condensates in a distinct manner. A. Representative Airyscan images of D4 cells co-expressing Halo-GRdim or Halo-GRtetra labeled with JF549 (green) and GFP-tagged nano-antibodies against each of the phosphorylated forms of Pol2 (magenta). Scale bar, 5 µm. B. Colocalization analysis of the condensates of the different GR-variants and Pol2 Ser5P (left panel, light colors tones) or Pol2 Ser2P (right panel, dark color tones) foci. The average CCF curve for each experimental condition (orange or green lines, respectively) and that expected for uncorrelated events (black line) are shown. C. Quantification of the nuclear area occupied by the condensates of GR-variants and colocalizing with Pol2 Ser5P (initiation) or Pol2 Ser2P (elongation) foci. Data is expressed as means ± SEM. D. Nuclear area occupied by receptor’s condensates colocalizing with Pol2 Ser5P (top) or Pol2 Ser2P (bottom) foci as a function of the total density of condensates of GR-variants. E. Relative intensity of the condensates of GR-variants colocalizing with Pol2 Ser5P or Pol2 Ser2P foci. Data is expressed as means ± SEM. Data information: Data sets are representative of at least three independent experiments. The number of cells (n) was n_Halo-GR/GFP-Pol2 Ser5P_ = 23, n_Halo-GR/GFP-Pol2 Ser2P_ = 14. n_Halo-GRdim/GFP-Pol2 Ser5P_ = 31, n_Halo-GRdim/GFP-Pol2 Ser2P_ = 27, n_Halo-GRtetra/GFP-Pol2 Ser5P_ = 25, n_Halo-GRtetra/GFP-Pol2 Ser2P_ = 25. Statistical analysis was performed by Man Whitney’ test. ns = not significant; * p < 0.05; ** p < 0.01 and *** p < 0.001.

Next, we analyzed the mean intensity of initiation, elongation, and total (active + inactive) condensates for each GR variant. Transcriptional active condensates of GRdim were dimmer than the total condensates (Fig. S4D) as observed for WT GR (Fig. 1D). In contrast, the mean brightness of GRtetra condensates was only slightly different to that measured for initiation and elongation condensates (Fig. S4D), indicating a similar number of GRtetra molecules in active and inactive condensates.

We also analyzed the fractional area of each nucleus occupied by these condensates (f_nucleus_, Fig. S1B) as this parameter provides clues on the total number of receptor condensates engaged in either transcription stage. The analysis of initiation condensates revealed higher f_nucleus_ values for GRtetra and GRdim compared to that found for the WT receptor (Fig. 4C). In fact, there seems to be a correlation between the total number of condensates linked to transcription initiation and the total density of condensates in the nucleus (Fig. 4D), suggesting that the number of GR condensates associated with initiation depends on the receptor’s capacity to form condensates. In addition, the mean intensity analysis (Fig. 4E) shows a correlation between the number of receptor molecules per initiation condensate and the transcriptional ability of the GR variants, as GRdim initiation condensates were dimmer than those formed by the WT receptor whereas GRtetra presented the highest brightness for initiation condensates.

If we now focus on elongation condensates, the fractional area of GRdim condensates showed a marked reduction compared to the WT receptor (Fig. 4C), also in line with the CCF observations (Fig. 4B). In contrast, the area of GRtetra and GR condensates associated with elongation was similar between each other. In addition, the total number of condensates linked to transcription elongation does not seem to correlate with the total density of condensates in the nucleus as observed for initiation (Fig. 4D). Finally, the analysis of the receptor condensates’ brightness revealed that elongation condensates also recruit more GRtetra than WT GR molecules as it was observed for initiation condensates. On the other hand, GRdim presented the lowest recruitment of all (Fig. 4E), having fewer condensates associated with elongation, which also concentrate less GRdim molecules, probably related to its poor transactivation capabilities.

Collectively, our data suggests that subpopulations of GR condensates participate in initiation and elongation. The number of initiation condensates depends on the receptor’s capacity to form condensates overall, with a correlation between the number of receptor molecules per initiation condensate and the transcriptional ability of the GR variants. In addition, the number of elongating condensates and receptor molecules per condensate seems to reflect the transcriptional ability of the GR variant.

## Discussion

Our comprehension of transcription has undergone remarkable transformations in the last few years [39]. One of the most astonishing observations is the intricate, multiphase architecture of the nucleus, wherein certain membrane-less, subnuclear compartments gather transcriptional players that interact via weak, multivalent interactions [40]. Initially, condensates were proposed to enhance transcription by concentrating several molecular actors thus increasing the probability of productive interactions [41]. However, it has been shown that accumulating these proteins in condensates does not necessarily lead to enhanced transcription [42-44]. Evidence points to a more complex scenario wherein transcriptional condensates with varying compositions and functions can coexist within a cell, potentially constituting an additional layer of temporal and spatial regulation of transcription [6]. Despite significant progress in this area, the role of these subnuclear structures in the regulation of transcription remains controversial.

The nuclear condensates formed by active steroid receptors constitute a good example of this kind of subnuclear structures with unknown roles [14], even though they were first documented decades ago [15]. In this work, we explored the potential role of GR condensates in transcription by using Airyscan microscopy, which allowed us to capture details hidden in standard confocal microscopy methods. We also employed specific genetically encoded nano-antibodies designed to label Pol2 Ser5P and Pol2 Ser2P, allowing us to visualize endogenous sites of active transcription initiation and elongation in live cells, respectively. By combining super-resolution microscopy and live cell imaging we provide evidence supporting the engagement of GR condensates in both stages of the transcription process.

Our spatial cross-correlation analyses showed spatial correlations between GR condensates and active Pol2 foci at a zero-distance shift. Although the colocalization analyses are limited by the Airyscan resolution, these observations suggest that even if they do not coexist in the same condensate or macromolecular structure, they must be closely associated in space. Moreover, GR condensate composition appears sensitive to perturbations in the molecular interactions occurring during different transcription stages. For example, the overexpression of Mediator subunit Med1 modulated the GR concentration at condensates. Consistently, blocking the switch of Pol2 from initiation to elongation promoted the dissociation of GR molecules from condensates, similar to what we observed at a tandem array of a GR reporter gene.

Transcription has been observed to occur in bursts, i.e., periods of active transcription resulting in numerous transcripts interspersed with inactive periods [45]; this phenomenon is likely the consequence of the transient, stochastic interactions between TFs, their cognate DNA targets, Pol2, and other transcriptional players [45]. Condensates might facilitate burst initiation as they provide a locally high concentration of key transcriptional players [46, 47]. The quantitative analysis of colocalization with the active forms of Pol2 is consistent with this paradigm as most GR condensates remain inactive within the 18.4 s window of the snapshot, while only a few condensates are involved in transcription initiation or elongation. It is tempting to speculate that, as a reflection of the status of its target genes, GR condensates mirror the periods of inactivity interspersed with periods of active transcription.

As the mean intensities of inactive, initiation, and elongating GR condensates are different, the transition between these states may be triggered or accompanied by changes in the relative composition of the condensates. In support of this idea, the intensity distribution of GR condensates spans over a large range of values, indicating a heterogeneous population of condensates as expected for these non-stoichiometric structures. In turn, the internal composition responds to interactions with transcriptional players as illustrated in experiments where Med1 was overexpressed, or transcriptional elongation was suppressed. Of note, the reduction of condensate’s mean intensity produced by Med1 overexpression is in line with the lower mean intensity of initiation condensates compared to inactive condensates. Consistent with our findings, it has been proposed that transcription progression involves a gradual modification in the condensate’s composition produced by the dissociation, incorporation or transformation of its molecules [48]. Importantly, although we are analyzing the intensity of fluorescently tagged GR at condensates, these structures also concentrate coactivators and other transcriptional players, not probed in our experiments. Last, it is important to emphasize that our results do not rule out the existence of a subpopulation of GR condensates which might not participate in transcription at all along their lifecycle.

Unexpectedly, our data points to a fraction of GR condensates colocalizing with transcription elongation foci, even though at a lower proportion than with initiation foci. Further work is needed to understand why a TF will remain close by during elongation, behaving differently than other molecules ‘purely’ related to transcription initiation, as we observed for the coactivator BRD4. At present, we cannot rule out that GR molecules are passively moving along these elongating condensates as there are no reports to the best of our knowledge of an active involvement of GR in elongation. Perhaps we are observing a small fraction of clustering at enhancers located within the gene body or additionally, GR could contribute to the weak-interactions network stabilizing the elongation condensates. Of note, Mediator plays a role in many stages of transcription, including initiation and elongation [49] and GR presents direct and indirect interactions with some of its subunits [32]. In addition, the androgen receptor, another member of the SRs family, interacts with the elongation factor pTEFb and enhances the efficiency of this process [50].

Noticeably, while we detected only ∼10 % of colocalized areas between GR and elongating condensates, DRB promoted the dissolution of ∼ 45 % of the receptor condensates. These seemingly contradictory results could be explained by technical limitations of our set up and/or confounding effects from inhibiting transcription. From a technical perspective, it is possible that many elongation foci are not detected in our two-color imaging experiments. Indeed, Pol2 clusters with as low as 5-10 molecules have been reported [34]. Additionally, the stoichiometry of labeling of the elongating polymerase depends on the expression level of the nano-antibody and might also be heterogeneous across the nuclear space due to the multivalency of CTDs [23]. Altogether, these limitations could result in an underestimation of elongating foci in Pol2 Ser2 images. Finally, disrupting transcription elongation produces changes in genome organization and affects the compartmentalization of several nuclear proteins [51]. In turn, these effects may also induce a reorganization of some components of GR condensates and trigger their partial dissolution. Further work is needed to determine whether GR condensates maintenance truly depends on the active transcription process, or whether these observations are also an indirect result of other effects produced by transcription inhibition.

To get insights into the functional role of GR condensates, we analyzed two mutants of the receptor with opposite transcriptional properties. The constitutively tetrameric GRtetra regulates a greater number of genes compared to the wild-type receptor [36], and GRdim remains a good chromatin binder but poorly activates the transcription of target genes [25]. Despite these dissimilar functional properties, both mutants form a large number of condensates after hormone activation, suggesting a poor correlation between condensate formation capability and transcriptional activity.

Interestingly, condensates formed by GRtetra and GRdim differ in several properties. First, their intensity distributions showed different behaviors to WT GR, wherein GRdim distribution shows a lower contribution of brighter condensates while GRtetra presents the opposite behavior. The latter is likely due to GRtetra’s enhanced multivalency that allows the interaction with many more molecular partners. Second, while the condensates formed by both mutants colocalize with initiation and elongation foci, they exhibit distinct characteristics. For example, the number of condensates engaged in transcription initiation seems to be proportional to the total density of condensates observed for each GR variant, suggesting that the number of initiation condensates is simply related to the ability of the receptor to form condensates. However, the number of receptors’ molecules recruited in each condensate (i.e. the condensate brightness) correlates with transcriptional activity as initiation condensates formed by GRdim were dimmer than those formed by the wild-type receptor, while GRtetra displayed the highest brightness. Finally, the engagement of each GR variant into elongating condensates appears to be linked to the receptor’s transcriptional activity, as GRdim formed fewer elongating condensates than the wild-type receptor and GRtetra. Additionally, the comparison of condensate’s mean intensities revealed that elongation condensates, similarly to initiation condensates, incorporate more GRtetra molecules than WT GR, with GRdim showing the lowest recruitment level. Thus, the ability of GR variants to participate in elongating condensates appears related to specific molecular determinants that define the number of receptor copies included in these condensates. As we discussed before, higher local concentrations of the receptor’s molecules possibly increase the probability of productive interactions leading to transcription events.

In conclusion, our data indicates a potential role for GR condensates in transcription. The enhanced transcriptional ability of GRtetra is consistent with higher participation in both initiation and elongation condensates, whereas the transcriptional impairment of GRdim is especially reflected in its lower concentration at initiation condensates, and reduced contribution to elongation condensates. This might be related to the defective behavior of GRdim in certain steps between initiation and elongation.

## Methods

### Cell culture and transfection

D4 (GR knock out) adenocarcinoma cell lines were previously described [25, 36]. Cells were cultured in DMEM high glucose (Gibco) supplemented with 10% FBS (Internegocios S.A.), 1% penicillin-streptomycin (Gibco), and 5 µg/ml tetracycline (Santa Cruz Biotechnology). Cells were maintained at 37 ºC under a humidified atmosphere with 5% CO_2_.

Transient transfections were performed with jetPRIME™ reagent (Polyplus, Sartorius) following the vendor’s instructions. Briefly, 2.5 × 10^5^ cells were grown on round coverslips and transfected with 1 µg of DNA for 4 h. Transfection medium was replaced with DMEM containing 5% charcoal-stripped FBS. Cells were incubated overnight with this medium prior to microscopy observation. The plasmids were pRNAP2 Ser2ph-mintbody (PB533-42B3mutAC2-sfGFP, Addgene #186777); pRNAP2 Ser5ph-mintbody (PB533-44B12m23-sfGFP, Addgene #186778); mCherry-BRD4 (Addgene #183939); pEGFP-GR [27]: Halo-Med1 (kindly provided by Joan Conaway, Stowers Institute, Kansas City, USA); pEGFP-GR407C; pEGFP-GRdim; pEGFP-GRtetra; Halo-GRdim and Halo-GRtetra were gifts from Dr. Gordon Hager (NIH, Bethesda, USA).

D4 cells expressing Halo-tagged proteins were incubated 40-60 min with the fluorescent dye JF549 (Janelia Farms, HHMI, USA) (50 nM) and then washed three times for 5 min before imaging.

### Sample preparation for imaging

Cells were grown on coverslips as described above and incubated with 100 nM Dexamethasone (Sigma–Aldrich) for 30 min at 37 ºC and 5% CO_2_ to induce GR nuclear translocation. Before imaging, the coverslips were mounted in a custom-made chamber designed for the microscope. DRB experiments were performed by incubating the cells with 100 µM of the drug (Sigma– Aldrich) for 30 min. This treatment was performed after the pre-incubation with dexamethasone described above.

### Microscopy

Confocal and Airyscan super-resolution images were acquired in a Zeiss LSM980 confocal microscope at the Weber Advanced Microscopy Center (FCEN, University of Buenos Aires) using a Plan-Apochromat 63x oil immersion objective (NA = 1.4). EGFP or GFP were excited with a diode laser of 488 nm. JF549-labeled Halo and mCherry were excited using a diode laser of 543 nm. The average power at the sample was ∼ 13 µW (488 nm) and ∼ 15 μW (543 nm). Fluorescence was registered with photomultipliers (confocal, EGFP 490-659 nm) or the AiryScan 2 detector using 495-550 nm (GFP or EGFP) and 574-720 nm (JF549-labeled Halo or mCherry) filtering. Two-color images were acquired in sequential mode. Airyscan images were registered with pixel size and dwell time of 43 nm and 16 µs, respectively. Microscopy measurements were run at 37 ºC and 5% CO_2_.

### Condensates/foci analysis

Airyscan super-resolution images were analyzed using ImageJ software (NIH, USA). The images of the nucleus were binarized through a thresholding procedure to quantify the nuclear area and to use it as a mask to quantify the nuclear mean fluorescence intensity (excluding nucleoli). Condensates and foci were identified as spots in the nucleus with intensities above a selected threshold (i.e., 1.4*nuclear mean intensity) and sizes > 6 pixels in the binary image.

The condensates/foci number, area and mean intensities were calculated using the ImageJ plugin “Analyze Particles” [52], and their intensity was divided by the nuclear intensity to obtain the relative intensity of these structures. Condensates density (δ_condensates_) was calculated as the ratio of the number of condensates to the nuclear area.

### Colocalization analysis

Spatial cross-correlation analysis was performed with the ImageJ JACoP plugin [53] using the Van Steensel procedure [15]. Briefly, the software enables quantitative cross-correlation analysis of two-color images by shifting (pixel by pixel) one of the images with respect to the other along the x-axis direction and calculating the Pearson coefficient at each shifting. The cross-correlation function (CCF) represents the Pearson coefficient as a function of the pixel shift (Δx) (Fig. S1A). For all experimental conditions, the CCF of the binary images of GR condensates and Pol2 foci was calculated up to Δx=20 pixels (0.86 µm) in the x and y coordinates (obtained after rotating 90° both images) and averaged. For each pair of images, we also estimated the CCF obtained for unrelated spatial distributions of Pol2 foci and GR condensates computing the CCF obtained by rotating one of the binary image 90° with respect to the other.

The overlapping area of GR condensates and Pol2 foci (i.e. colocalization area, A_col_) was quantified by multiplying their binary images and adding all the colocalizing pixels (Fig. S1B). The relative area of GR condensates colocalizing with those of Pol2 was calculated as A_col_/ A_GR_, where A_GR_ is the total area of GR condensates, calculated as the sum of pixels in the binary image of GR condensates. The nuclear coverage of the receptor co-localizing with the Pol2 forms was calculated as the quotient between the corresponding A_col_ and the nuclear area (A_nucleus_).

### Statistical analyses

Results were expressed as means ± SEM of at least three replicates. Statistical significance for mean pairwise comparisons was performed using Student’s t-test. Before the analysis, data were tested for normality and homogeneity of variances using Shapiro-Wilk and Levene tests, respectively. Data sets that did not exhibit a normal distribution were analyzed using the Mann-Whitney test. If variances were not equal, the unpaired t-test with Welch’s correction was employed. Differences were considered as significant at p <□0.05.

Statistical analyses were performed with GraphPad Prism software.

## Acknowledgments

The authors thank Victoria Repetto for technical assistance at the Weber’s Advance Microscopy Center. This research was partially supported by ANPCyT (PICT 2020-00818 to V.L., PICT PRH 2018-0573 to D.M.P.), Universidad de Buenos Aires (UBACyT 20020190100101BA to V.L.) but PICT funding disbursements were suspended following the Argentina government’s decision to cut financial support for scientific research across the country.

