## Supplemental Figures for "Hallmarks of glucocorticoid receptor condensates involvement in transcription regulation"

### Supplementary information

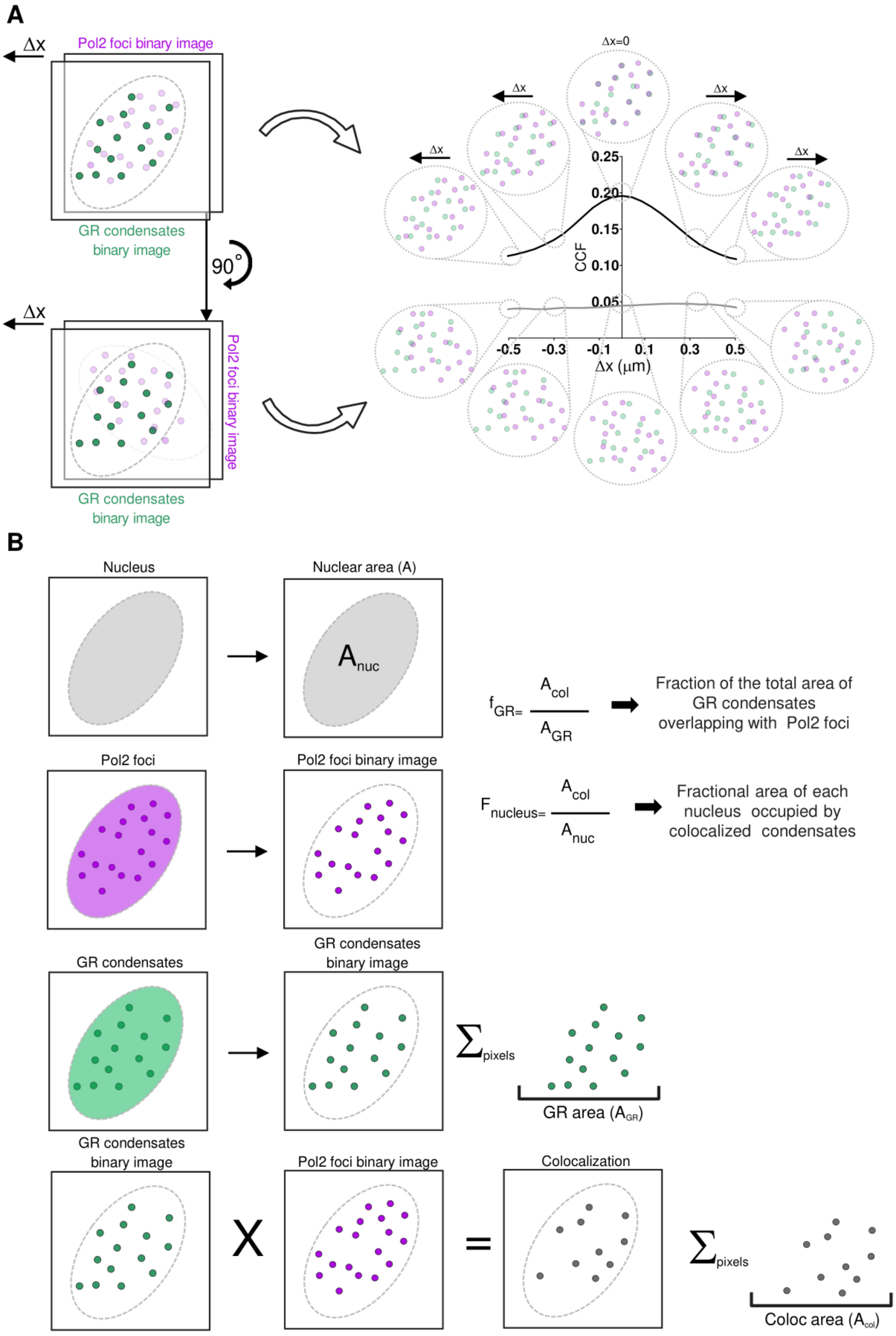

**S1. Schematic representation of the colocalization analysis between GR condensates and Pol2 foci.** A. Scheme showing the colocalization analysis performed using the ImageJ JACoP plugin. Microscopy images are processed to obtain the binary images for GR condensates (green) and Pol2 foci (magenta) (left panel). The Pol2 binary image is then shifted pixel by pixel (Δx) along the x-axis direction with respect to the GR binary image, computing the Pearson coefficient at each iteration. The same procedure was performed by rotating the Pol binary image 90° to obtain the result for unrelated events. The cross-correlation function (CCF) represents the Pearson coefficient as a function of the pixel shift (Δx), (right panel). The average CCF expected for correlated (i.e. colocalization) and uncorrelated (i.e. no colocalization) distributions are shown in black and grey, respectively. B. Diagram showing the quantification of the colocalizing area of GR condensates and Pol2 foci within the nucleus (dotted line). Microscopy images are segmented to obtain binary images of the cell nucleus (grey), GR condensates (green) and Pol2 foci (magenta) and compute their respective area. To calculate the total overlapping area between GR condensates and Pol2 foci ($A_{col}$), their binary images are multiplied and positive pixels of the colocalization image are added up to calculate the colocalizing area. This value is divided by the total area of GR condensates ($A_{GR}$) to estimate the relative area of the receptor´s condensates colocalizing with Pol2 foci ($A_{col}/A_{GR}$). Nuclear coverage of GR condensates colocalized with the active Pol2 foci was calculated as the ratio of the $A_{col}$ to the nuclear area ($A_{nucleus}$).

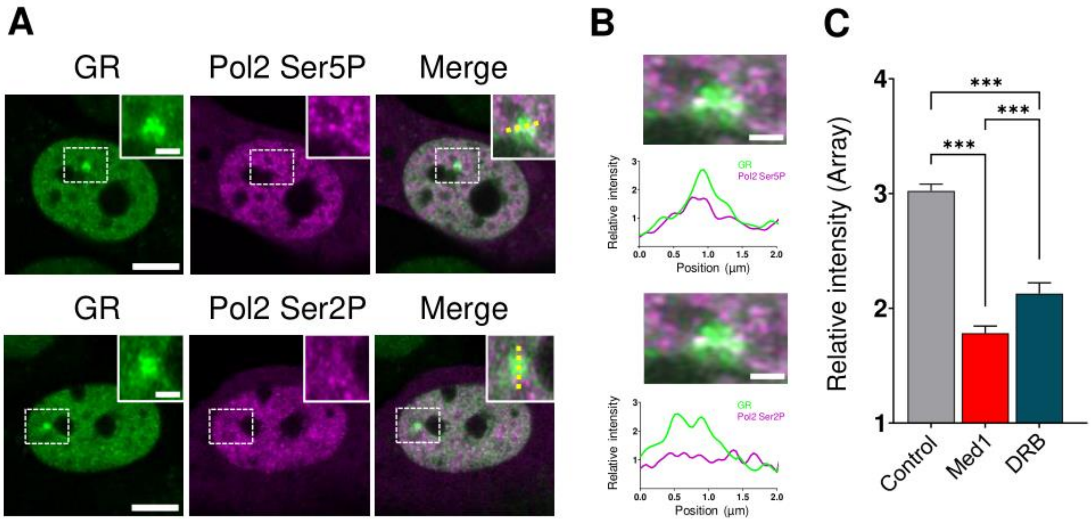


**S2. Relative distributions of active Pol2 foci and the MMTV-array.** A. Representative Airyscan images of D4 Halo-GR cells labeled with JF549 (green) and transiently expressing GFP-tagged nano-antibodies against Pol2 Ser5P) or Pol2 Ser2P (magenta). Scale bar, 5 µm. Zoom-in images of the region are delimited by the white squares including the MMTV-array. Scale bar, 1 µm. B. Zoom-in images of the region delimited by the white squares including the MMTV-array (top). Scale bar, 1 µm. Fluorescence intensity profile of GR (green) and Pol2 Ser5P or Pol2 Ser2P (magenta) registered along a line (yellow dotted line in panel A) that intercepts the MMTV-array (bottom). C. Relative intensity of MMTV-array in Dex-stimulated D4 cells expressing EGFP-GR (control), co-expressing EGFP-GR and Halo-Med1 labeled with JF549 (Med1) or after DRB incubation (DRB). Data is expressed as means ± SEM. Data information: Data sets are representative of at least three independent experiments. The number of cells (n) was: (C) $n_{EGFP-GR}$ = 43, $n_{EGFP-GR/Halo-Med1}$ = 19 and $n_{EGFP-GR+DRB}$ = 39. Statistical analysis was performed by Man Whitney' test. ns = not significant; * $p < 0.05$; ** $p < 0.01$ and *** $p < 0.001$.

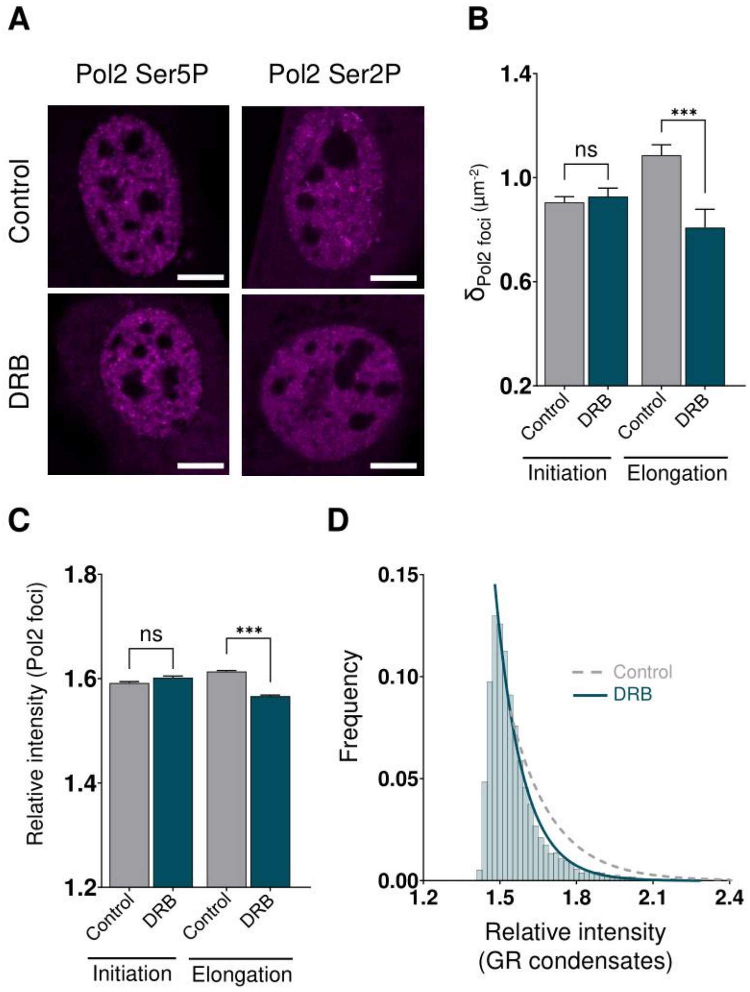


**S3. Pharmacological inhibition of transcription elongation affects Pol2 elongation foci but not Pol2 initiation foci.** A. Representative Airyscan images of D4 cells transiently expressing GFP-tagged nano-antibodies against Pol2 Ser5P (left) or Pol2 Ser2P (right) acquired in control condition and after incubation with DRB. Scale bar, 5 µm. B. Density of Pol2 Ser5P (initiation) and Pol2 Ser2P (elongation) foci quantified in control and DRB-treated cells. Data is expressed as means ± SEM. C. Relative intensity of Pol2 Ser5P and Pol2 Ser2P foci registered in control and DRB-treated cells. Data is expressed as means ± SEM. D. Intensity distribution of GR condensates in DRB-treated cells. The data was fitted with an exponential-decay function (green line) obtaining the parameters reported in Table S1. To facilitate comparison, the fitting curve obtained for GR in control condition is also shown (grey dotted line). Data information: Data sets are representative of at least three independent experiments. The number of cells (n) was (B-C) $n_{GFP\text{-}Pol2\ Ser5P\ (Control)} = 26$, $n_{GFP\text{-}Pol2\ Ser5P\ (DRB)} = 15$, $n_{GFP\text{-}Pol2\ Ser2P\ (Control)} = 14$, $n_{GFP\text{-}Pol2\ Ser2P\ (DRB)} = 10$; (D) $n_{EGFP\text{-}GR\ (control)} = 46$, $n_{EGFP\text{-}GR\ (DRB)} = 70$, $n_{EGFP\text{-}GR\ (Wash)} = 29$. Statistical analysis was performed by Student's t-test or Man Whitney' test. ns = not significant; * $p < 0.05$; ** $p < 0.01$ and *** $p < 0.001$.

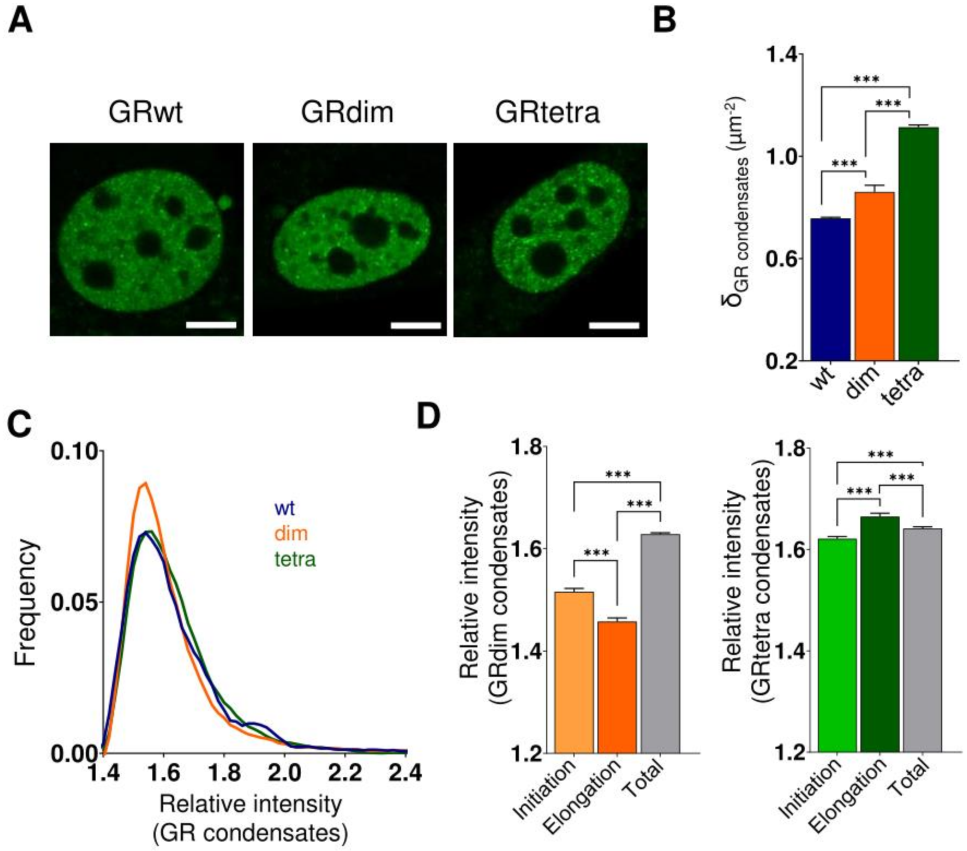


**S4. Distinct properties of condensates formed by different GR mutants.** A. Representative Airyscan images of D4 cells transiently transfected with EGFP-GRwt, EGFP-GRdim or EGFP-GRtetra. Scale bar, 5 µm. B. Nuclear density of condensates of the different GR variants. Data is expressed as means ± SEM. C. Intensity distribution of condensates formed by different GR mutants. D. Relative intensity of the total GRdim (left) or GRtetra (right) condensates (total) and those colocalizing with Pol2 Ser5P (initiation) or Pol2 Ser2P (elongation) foci. Data is expressed as means ± SEM. Data information: Data sets are representative of at least three independent experiments. The number of cells (n) was: (B-C) $n_{EGFP\text{-}GRwt}$ = 46, $n_{EGFP\text{-}GRdim}$ = 35, $n_{EGFP\text{-}GRtetra}$ = 30; (D) $n_{Halo\text{-}GR/GFP\text{-}Pol2\ Ser5P}$ = 23, $n_{Halo\text{-}GR/GFP\text{-}Pol2\ Ser2P}$ = 14. $n_{Halo\text{-}GRdim/GFP\text{-}Pol2\ Ser5P}$ = 31, $n_{Halo\text{-}GRdim/GFP\text{-}Pol2\ Ser2P}$ = 27, $n_{Halo\text{-}GRtetra/GFP\text{-}Pol2\ Ser5P}$ = 25, $n_{Halo\text{-}GRtetra/GFP\text{-}Pol2\ Ser2P}$ = 25. Statistical analysis was performed by unpaired t test with Welch's correction or Man Whitney' test. ns = not significant; * $p < 0.05$; ** $p < 0.01$ and *** $p < 0.001$.

**Table S1. Parameters of the exponential fit on the intensities of the GR condensates**

| Condition | Mean ± SEM | σ (†) ± SE | $N_{Condensates}$ / $N_{Cell}$ |
|---|---|---|---|
| WT GR | 1.645 ± 0.005 | 0.165 ± 0.007 | 1949 / 25 |
| GR407C | 1.609 ± 0.002 (*) | 0.118 ± 0.004 | 2836 / 40 |
| WT GR + Med1 | 1.541 ± 0.002 (*) | 0.087 ± 0.006 | 1471 / 22 |
| WT GR + DRB | 1.561 ± 0.002 (*) | 0.108 ± 0.004 | 5689 / 70 |
| GRdim | 1.628 ± 0.003 (*) | 0.132 ± 0.003 | 5174 / 37 |
| GRtetra | 1.641 ± 0.003 | 0.176 ± 0.009 | 2665 / 25 |

Asterisks indicate significant differences with WT GR ($p < 0.05$)

(†) σ= decay constant obtained by fitting to the right branch of the intensity distribution of condensates an exponential decay function ($A * e^{-x/\sigma}$), where x is the relativity intensity of GR condensates.
